# Development and characterization of nanobodies specifically targeting the oncogenic Phosphatase of Regenerating Liver-3 (PRL-3)

**DOI:** 10.1101/2020.10.02.311787

**Authors:** Caroline N. Smith, Kyle Kihn, Zachary A. Williamson, K. Martin Chow, Louis B. Hersh, Konstantin V. Korotkov, Daniel Deredge, Jessica S. Blackburn

## Abstract

Phosphatase of Regenerating Liver-3 (PRL-3) is associated with cancer progression and metastasis in various solid tumors and leukemias. The mechanisms that drive PRL-3’s oncogenic functions are not well understood, in part due to a lack of research tools available to study this protein. In particular, small molecules do not exhibit binding specificity for PRL-3 over highly homologous family members PRL-1 and PRL-2, and antibodies directed against PRL-3 are limited by assay type. We have begun to address these issues by developing alpaca-derived single domain antibodies, or nanobodies, targeting PRL-3 with a K_D_ of 30-300 nM and no activity towards PRL-1 and PRL-2. Hydrogen deuterium exchange mass spectrometry (HDX-MS) and co-immunoprecipitation with a known PRL-3 substrate showed the nanobodies bind PRL-3 outside of the active site, meaning they can be used to study PRL-3 interaction with binding partners. The nanobodies were also specific to PRL-3 over other PRLs in immunoprecipitation and immunofluorescence experiments in human cancer cells that overexpressed the PRL family. We found that N-terminal tags on PRL-3, such as GFP and FLAG, changed PRL-3 localization compared to untagged protein, indicating that the nanobodies may provide new insights into PRL-3 trafficking and function. The anti-PRL-3 nanobodies represent an important expansion of the research tools available to study PRL-3 function and can be used to define the role of PRL-3 in cancer progression.

## Introduction

The Protein Tyrosine Phosphatase 4A (PTP4A) family of three proteins, also known as Phosphatases of Regenerating Liver (PRL-1, PRL-2, and PRL-3), act as oncogenes in multiple cancer types. PRL-3, in particular, has been identified as a potential biomarker of cancer progression and metastasis in colorectal (1,2), gastric (3), ovarian (4), breast (5), brain (6), and prostate (7) cancers, as well as melanoma (8,9), and leukemias (10,11). Experimental evidence indicates that PRL-3 expression increases proliferation, migration, and invasion of cancer cells *in vitro* (12–14) and enhances tumor growth and metastasis in mouse models (2,15). In contrast, PRL-3 knockdown significantly suppresses tumor formation and spread *in vivo* (16). PRL-3 is well-established in inhibiting apoptosis, promoting epithelial to mesenchymal transition (EMT), and inducing migration in cancer cells. However, there are many unanswered questions regarding how PRL-3 drives these oncogenic phenotypes. PRL-3 has been characterized as both a functioning phosphatase involved in a variety of signaling pathways, including PTEN/PI3K/AKT, Src, STAT3/5, MAPK, and Rho GTPase, among others (17,18), as well as a pseudo-phosphatase that regulates magnesium transport via interaction with CNNM proteins (19,20). PRL-3 is mainly localized at the cell membrane (21,22) but is also found in the cytoplasm (8) and nucleus (23), where its substrates and functions are largely unknown.

Despite over two decades of research on PRL-3, there are many open questions regarding this protein, including its physiologic functions, trafficking, regulation, and *in vivo* substrates/binding partners. These unknowns are partly due to a lack of research tools available to study this protein. Developing small-molecule inhibitors specific to PRL-3 has been difficult, as PRL proteins are ~80% homologous, and the PRL catalytic binding pocket is both shallow and hydrophobic (24). Currently, the most frequently used PRL inhibitors are the PRL-3 Inhibitor I (Sigma P0108), Analog 3 (25), and thienopyridone (26), which appear to inactivate the PRL family via a redox reaction instead of directly binding with the protein’s active site. JMS-053, a thienopyridone derivative, is currently the only PRL inhibitor suitable for *in vivo* use and binds to PRL-3 in computational docking models. However, this has yet to be validated in molecular assays (27,28). Developing antibodies specific for PRL-3 has also proven difficult, with most antibodies lacking specificity towards PRL-3 over other PRL proteins. The only antibody validated across multiple research groups as specific for PRL-3 over PRL-1 and PRL-2 (26,29) was raised against the linear form of PRL-3 and therefore cannot be used for studies involving PRL-3 in its native structure. A humanized monoclonal antibody, PRL-3-zumab, binds specifically to PRL-3 and has anti-cancer effects *in vivo* (30). The authors of this work predicted that PRL-3 is presented on the cell surface through exosomal secretion, thereby allowing for the binding of PRL-3-zumab. This event stimulates Fc-receptor-dependent interactions between PRL-3 positive cells and host immune effectors, activating classical antibody-mediated tumor clearance pathways leading to tumor cell death (30). While PRL-3-zumab is currently in phase 2 clinical trial in Singapore (NCT04118114) for gastric and hepatocellular carcinomas and the United States (NCT04452955) for solid tumors, this antibody is not currently commercially available, which limits its research use. Overall, tools to study PRL-3 are lacking.

Single domain antibodies, or nanobodies, have recently emerged as important research tools and are likely to become useful therapeutics in various diseases, including cancer (31,32). Nanobodies were discovered in dromedaries, such as camels, llamas, and alpacas. These animals produce both antibodies with a typical structure and those with an atypical structure that lack light chains but have a similar variable region (VHH region) compared to conventional antibodies (33). The lack of light chains causes the formation of a longer complementary determining region-(CDR)3 with a secondary disulfide bond to stabilize the nanobody structure (34). This stabilization permits the formation of convex shapes, allowing nanobodies to reach narrow, concave binding and activation sites on proteins that normal antibodies cannot (35). Other advantages of nanobodies include their small size of ~15 kD, stability under stringent conditions, lack of immunogenicity, and high specificity and affinity for their antigens (33).

These same properties make nanobodies advantageous as therapeutics in cancer applications (36). The small size of nanobodies enables deep penetration into tumor tissue and the ability to cross the blood-brain barrier (37) while maintaining low off-target effects (38). Nanobodies can also withstand high temperatures, elevated pressure, non-physiological pH, and denaturants, making them ideal research tools and functional therapeutic molecules (36). In 2019, the first nanobody therapeutic, Caplacizumab or Cablivi, was approved by the FDA to aid in accelerating platelet aggregation in acquired thrombotic thrombocytopenic purpura (aTTP), a disease that causes small blood clots throughout the body. There are currently over 12 ongoing clinical trials examining the efficacy of nanobodies across a range of cancer types (36). Over the last ten years, there has been consistent curiosity in what we can learn about target proteins using nanobodies (36,39,40).

Here, we describe the development and characterization of the first nanobodies against PRL-3. We show that anti-PRL-3 nanobodies can be used in biochemical assays such as ELISA, where they interact with PRL-3 without binding to PRL-1 or PRL2. The binding affinity between nanobodies and PRL-3 was in the nanomolar range (41), similar to many antibodies therapeutics currently on the market. We performed epitope mapping using hydrogen-deuterium exchange mass spectrometry (HDX-MS) to characterize nanobody interaction with PRL-3 and found that nanobodies bound away from the PRL-3 active site. Nanobodies did not interfere with PRL-3 phosphatase activity and allowed for immunoprecipitation of purified PRL-3 protein with a CNNM peptide substrate. Anti-PRL-3 nanobodies successfully immunoprecipitated PRL-3 protein from cell lysates and co-localized with PRL-3 in immunofluorescence assays in colorectal cancer cells, with no interaction with PRL-1 or PRL-2. In total, these anti-PRL-3 nanobodies are among the first tools in the PRL field that can specifically detect native PRL-3 protein in a range of biochemical and cellular assays. PRL-3 nanobodies will be helpful in defining the normal and oncogenic functions of PRL-3 and may aid in the development of novel therapeutics targeted to this protein.

## Results

### Alpaca derived anti-PRL-3 nanobodies exhibit varying amino acid sequences as well as abundance following phage display

Human recombinant PRL-3 protein was injected into alpacas, and B-lymphocytes expressing potential anti-PRL-3 nanobodies were harvested six weeks later and developed into a cDNA library, as diagramed in Figure 1A. Bacteriophage display panning of the library against recombinant PRL-3 and subsequent sequencing of the enriched clones identified 32 potential nanobodies. Of these, only 16 nanobodies contained a complete N-terminal PelB sequence for packaging the protein in the bacterial periplasm for protein purification, a C-terminal 6XHis-tag for immobilized metal affinity chromatography purification, a stop codon, and were without any undetermined amino acids.

**Figure 1.**
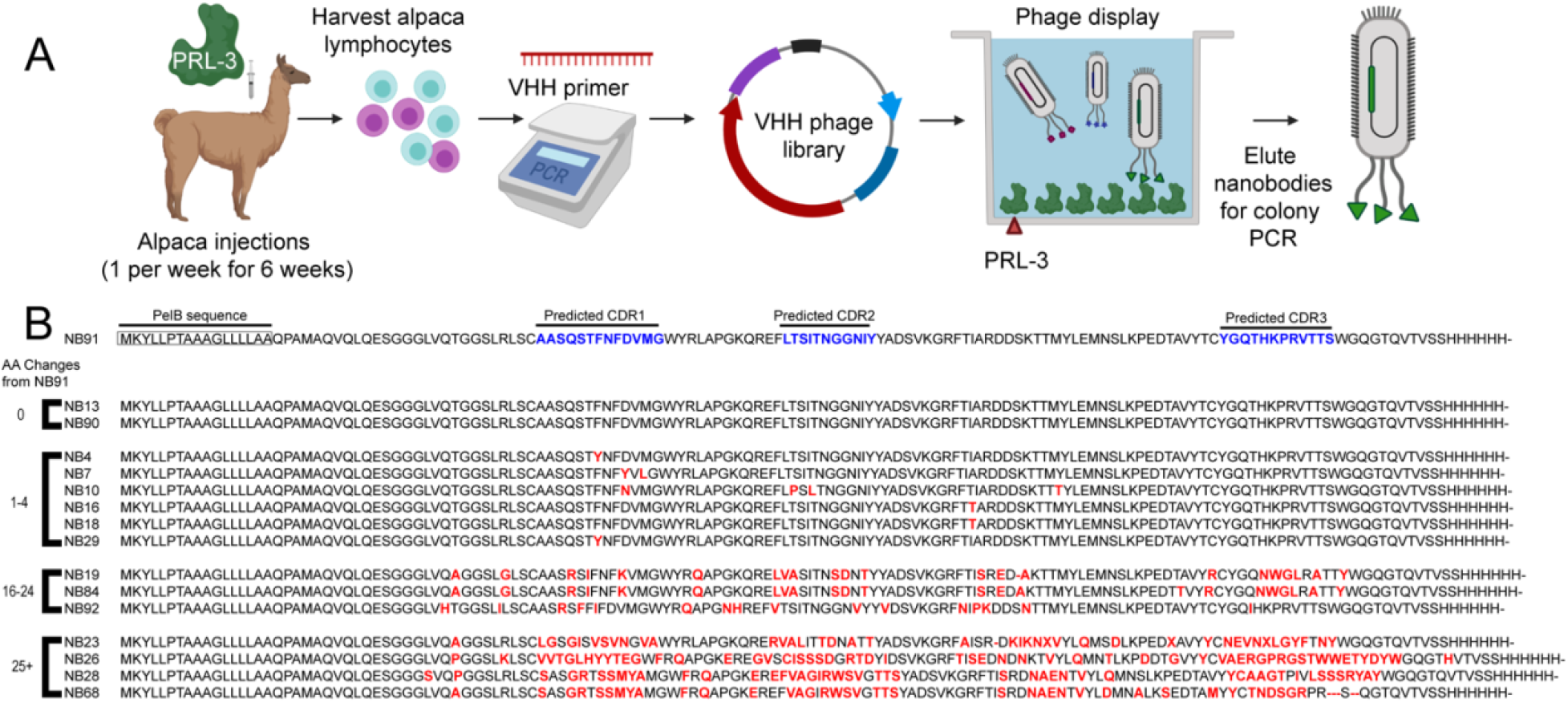
Isolation of PRL-3 specific nanobodies from alpacas. (A) Schematic demonstrating the process completed from initial recombinant PRL-3 injection in alpacas to colony PCR to sequence potential anti-PRL-3 nanobodies (Created with BioRender.com). (B) Amino acid sequence for 16 nanobodies used in this study. Nanobodies are grouped based on amino acid similarity to NB91. Each group of nanobodies is either the same sequence as NB91, differs by 1-4, 10-25, or 25+ amino acids (red). Complimentary determining regions (CDs, blue in nanobody 91) were predicted using ABodyBuilder.

Alignment of the anti-PRL-3 nanobody amino acid sequences showed nanobodies 91, 90, and 13 were 100% identical (Figure 1B). This sequence was the most recurrent nanobody sequence identified; therefore, we utilized nanobody 91 as our standard anti-PRL-3 nanobody throughout this study. The complementary-determining regions of this nanobody were predicted using ABodyBuilder (42). Nanobody sequences were clustered based on the number of amino acid alterations or insertions compared to nanobody 91. These include four groups containing 0, 1-4, 10-20, and 25+ amino acid changes compared to nanobody 91.

### Anti-PRL-3 nanobodies are specific for PRL-3 over other PRL family members in protein assays

PRL-1 and PRL-2 have 79% and 76% amino acid sequence homology to PRL-3 (24), making identifying specific small molecules and antibodies challenging. We used an indirect ELISA method to test the specificity of the anti-PRL-3 nanobodies towards PRL-3 over other PRL family members. We found that 11 of 16 nanobodies had a significantly greater affinity for PRL-3 over PRL-1 and PRL-2 (Figure 2A). Even under saturating conditions, most anti-PRL-3 nanobodies lacked any binding to PRL-1 or PRL-2 protein (Figure 2B). As expected, nanobodies with similar amino acid sequences had comparable binding to PRL-3. Five nanobodies had reduced binding to PRL-3; we attributed this to poor expression in *E. coli* (nanobodies 7, 68, and 92) or low nanobody affinity for PRL-3 (nanobodies 23 and 28). Our further studies focus on seven nanobodies (4, 10, 16, 19, 26, 84, and 91) with a strong affinity for PRL-3 and unique amino acid sequences compared to nanobody 91.

**Figure 2.**
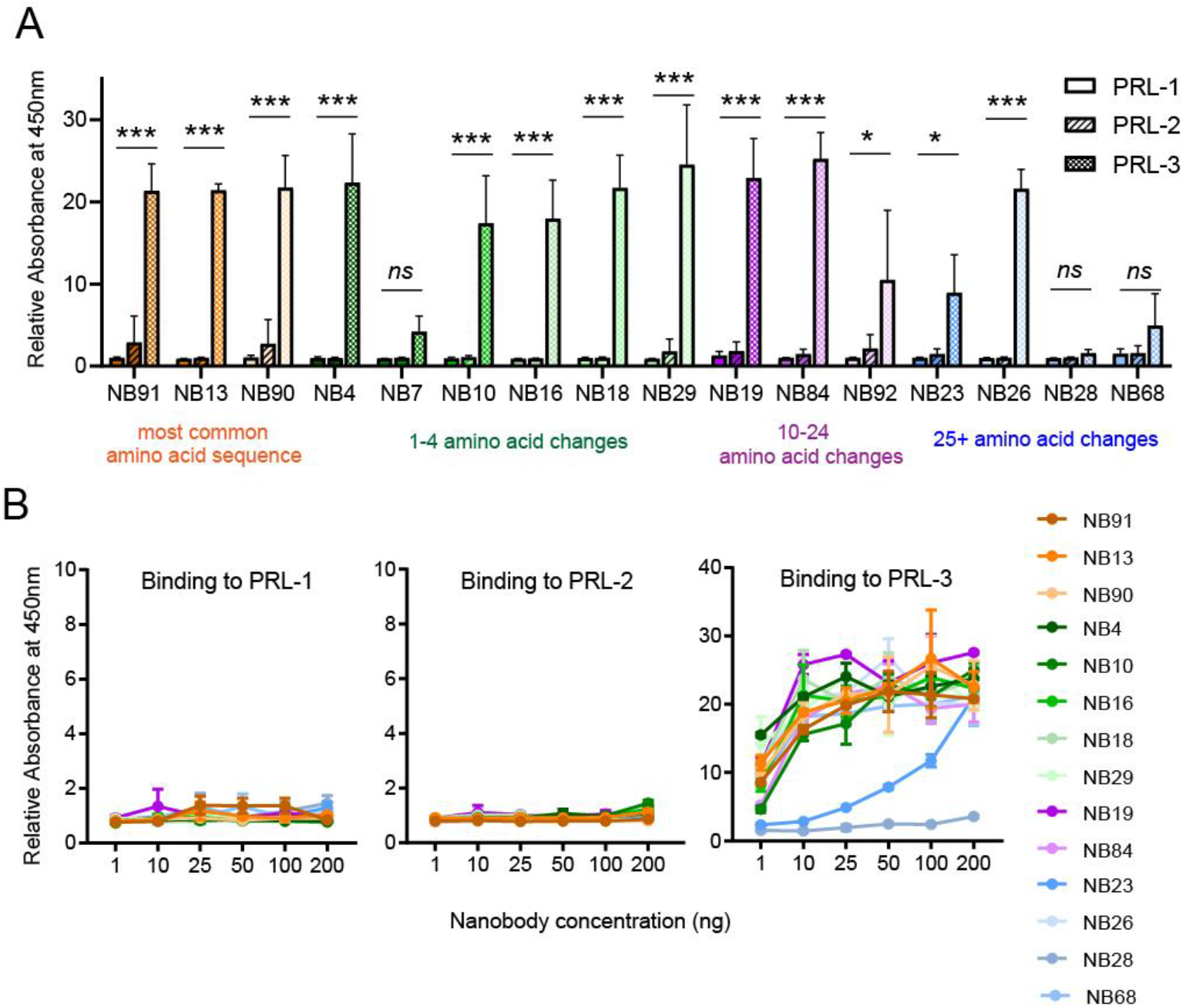
Nanobodies are specific for PRL-3 over the other PRL family members. (A) Binding of each histidine-tagged nanobody (NB, at a concentration of 100 ng) to 100 ng PRL-1, PRL-2, or PRL-3 in 96-well plates. (B) The binding of each nanobody at the concentrations indicated to 100 ng of each PRL was measured by indirect ELISA. Data are the absorbance at 450 nm after NB/PRL wells were washed and probed with His-HRP conjugated antibody. All assays were completed with two technical replicates and repeated in two biological replicates. Error bars represent standard deviation. ns = not significant, \**p* < 0.05, \*\*\*\**p* < 0.0001 by two-way ANOVA with Sidak’s multiple comparisons test. The number of amino acid changes compared to the most common anti-PRL-3 nanobody sequence is indicated by color-coding.

### Anti-PRL-3 nanobodies have a binding affinity for PRL-3 in the nanomolar range

We determined each nanobody’s dissociation constant (K_D_) for PRL-3 using Biolayer Interferometry (BLI). A smaller dissociation constant correlates to a higher affinity constant and indicates a stronger interaction between PRL-3 and nanobody. We calculated the global K_D_ from six increasing concentrations of PRL-3 for each nanobody (Tables S1). Nanobodies 19 and 26 have the greatest affinity for PRL-3, at 98.4 nM and 28.9 nM, respectively (Table 1). On average, commercially available and FDA-approved antibodies have affinities ranging from 10^-5^ to 10^-11^ M to their targets; our anti-PRL-3 nanobodies are well within this range.

**Table 1.**
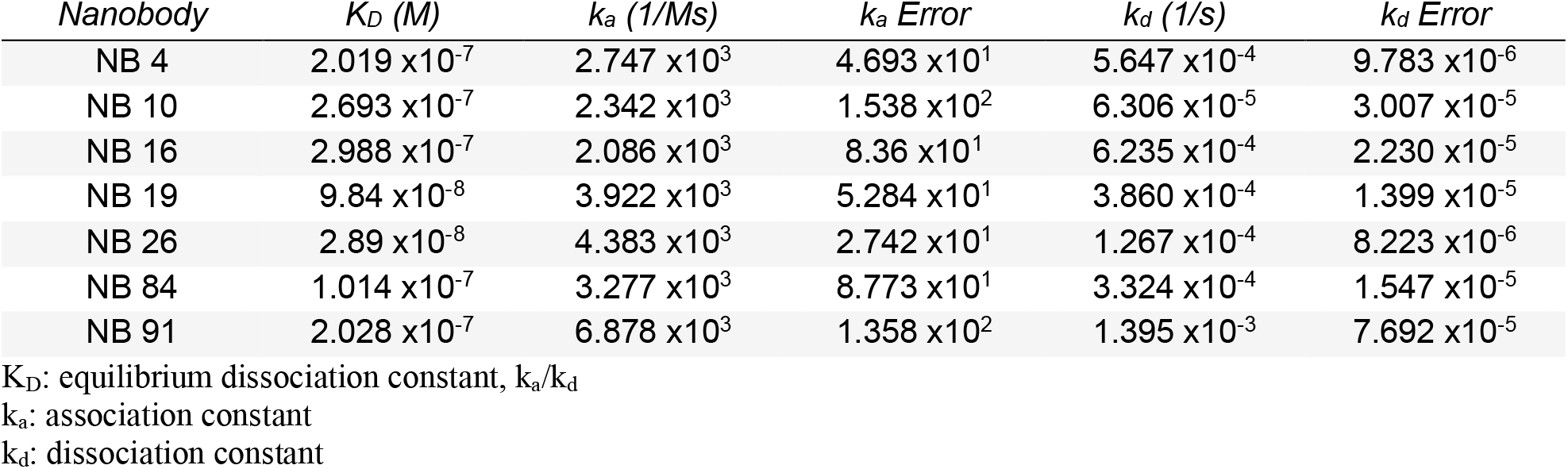
Kinetics of nanobody binding to PRL-3.

| *Nanobody* | $K_D$ (M) | $k_a$ (1/Ms) | $k_a$ Error | $k_d$ (1/s) | $k_d$ Error |
| --- | --- | --- | --- | --- | --- |
| NB 4 | $2.019 \times 10^{-7}$ | $2.747 \times 10^3$ | $4.693 \times 10^1$ | $5.647 \times 10^{-4}$ | $9.783 \times 10^{-6}$ |
| NB 10 | $2.693 \times 10^{-7}$ | $2.342 \times 10^3$ | $1.538 \times 10^2$ | $6.306 \times 10^{-5}$ | $3.007 \times 10^{-5}$ |
| NB 16 | $2.988 \times 10^{-7}$ | $2.086 \times 10^3$ | $8.36 \times 10^1$ | $6.235 \times 10^{-4}$ | $2.230 \times 10^{-5}$ |
| NB 19 | $9.84 \times 10^{-8}$ | $3.922 \times 10^3$ | $5.284 \times 10^1$ | $3.860 \times 10^{-4}$ | $1.399 \times 10^{-5}$ |
| NB 26 | $2.89 \times 10^{-8}$ | $4.383 \times 10^3$ | $2.742 \times 10^1$ | $1.267 \times 10^{-4}$ | $8.223 \times 10^{-6}$ |
| NB 84 | $1.014 \times 10^{-7}$ | $3.277 \times 10^3$ | $8.773 \times 10^1$ | $3.324 \times 10^{-4}$ | $1.547 \times 10^{-5}$ |
| NB 91 | $2.028 \times 10^{-7}$ | $6.878 \times 10^3$ | $1.358 \times 10^2$ | $1.395 \times 10^{-3}$ | $7.692 \times 10^{-5}$ |

$K_D$ : equilibrium dissociation constant, $k_a/k_d$
$k_a$ : association constant
$k_d$ : dissociation constant

### HDX-MS defines two regions of interaction and stabilization between PRL-3 and three nanobodies

We used Hydrogen Deuterium Exchange Mass Spectrometry (HDX-MS) to define the potential binding sites between nanobodies 19, 26, and 91 and PRL-3. HDX-MS probes the structure of a protein complex by monitoring the exchange of backbone amide hydrogen atoms with solvent deuterium atoms upon exposure to deuterated solvent. We compared the deuterium uptake of apo-PRL-3 to PRL-3 complexed with either nanobody 91, 19, or 26. We found that nanobody interaction with PRL-3 protected PRL-3 from deuteration in two regions, shown in blue in Figures 3A, S1, and S2. We unexpectedly found an increase in deuteration, or deprotection, of up to 30% in one region of PRL-3 after nanobody binding (shown in red), which may indicate a slight conformational change in PRL-3 caused by nanobody binding, compared to apo-PRL-3. One caveat to this experiment was our inability to determine structural interactions at the PRL-3 active site, between amino acids 72 to 104, due to a lack of sequence coverage. Overall, nanobody 91 deprotected PRL-3 from residues 13-19 and protected at amino acids 56-79 and 132-146 (Figure 3B). Nanobody 19 and 26 shared similar alterations in deuterium uptake compared to nanobody 91, with one deprotection event and two protection events in similar regions (Figures S1-S2). Through this pattern of protection and deprotection, we conclude that all three nanobodies appear to bind very similar regions of PRL-3 and would likely have similar effects on overall protein structure and stability.

**Figure 3.**
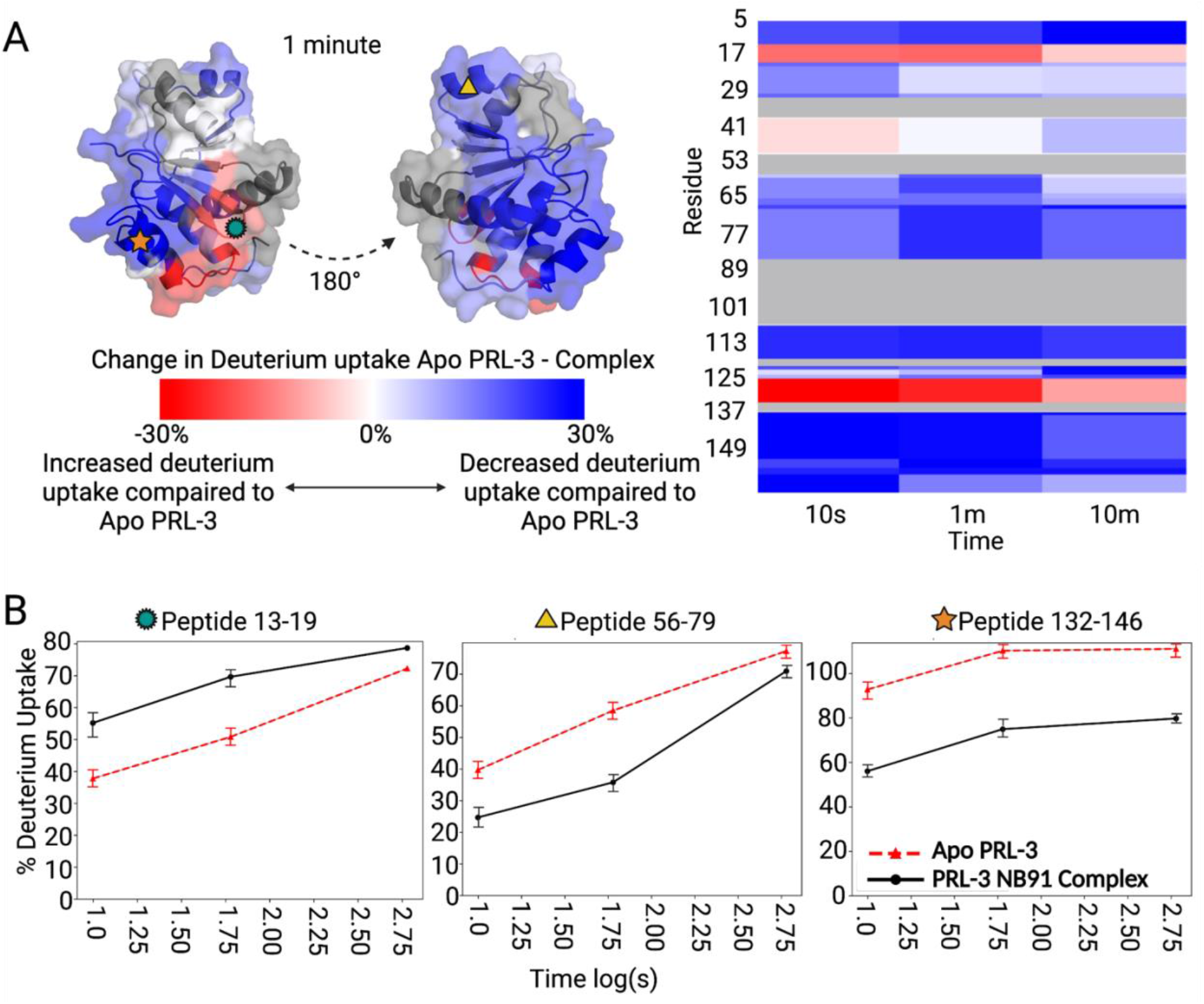
Hydrogen Deuterium Exchange Mass Spectrometry defines Nanobody 91 binding sites with PRL-3. (A) PRL-3 in complex with nanobody 91 shows regions of both increased (red) and decreased (blue) deuterium uptake, compared to apo-PRL-3. Heatmap indicates approximately 70% sequence coverage by mass spectrometry; gray areas represent portions of PRL-3 where data for deuterium exchange was not recovered. (B) Peptide 13-19 showed PRL-3 deprotected following nanobody binding, while peptides 56-79 and 132-146 showed decreases in deuterium uptake, reflecting more protection by nanobody 91 on PRL-3 in these regions.

### Anti-PRL-3 nanobodies do not inhibit PRL-3 phosphatase activity or substrate binding

The HDX-MS data indicate that the nanobodies are bound outside the PRL-3 active site; we wanted to experimentally confirm that PRL-3 activity was not negatively impacted by nanobody binding. We first assessed the impact of nanobody binding on the ability of PRL-3 to dephosphorylate a generic substrate, 6,8-Difluoro-4-Methylumbelliferyl Phosphate (diFMUP). We found no significant difference in PRL-3 activity between apo-PRL-3 and PRL-3:Nanobody complexes (Figures 4A, S3), suggesting that the nanobodies do not negatively impact the phosphatase activity of PRL-3.

**Figure 4.**
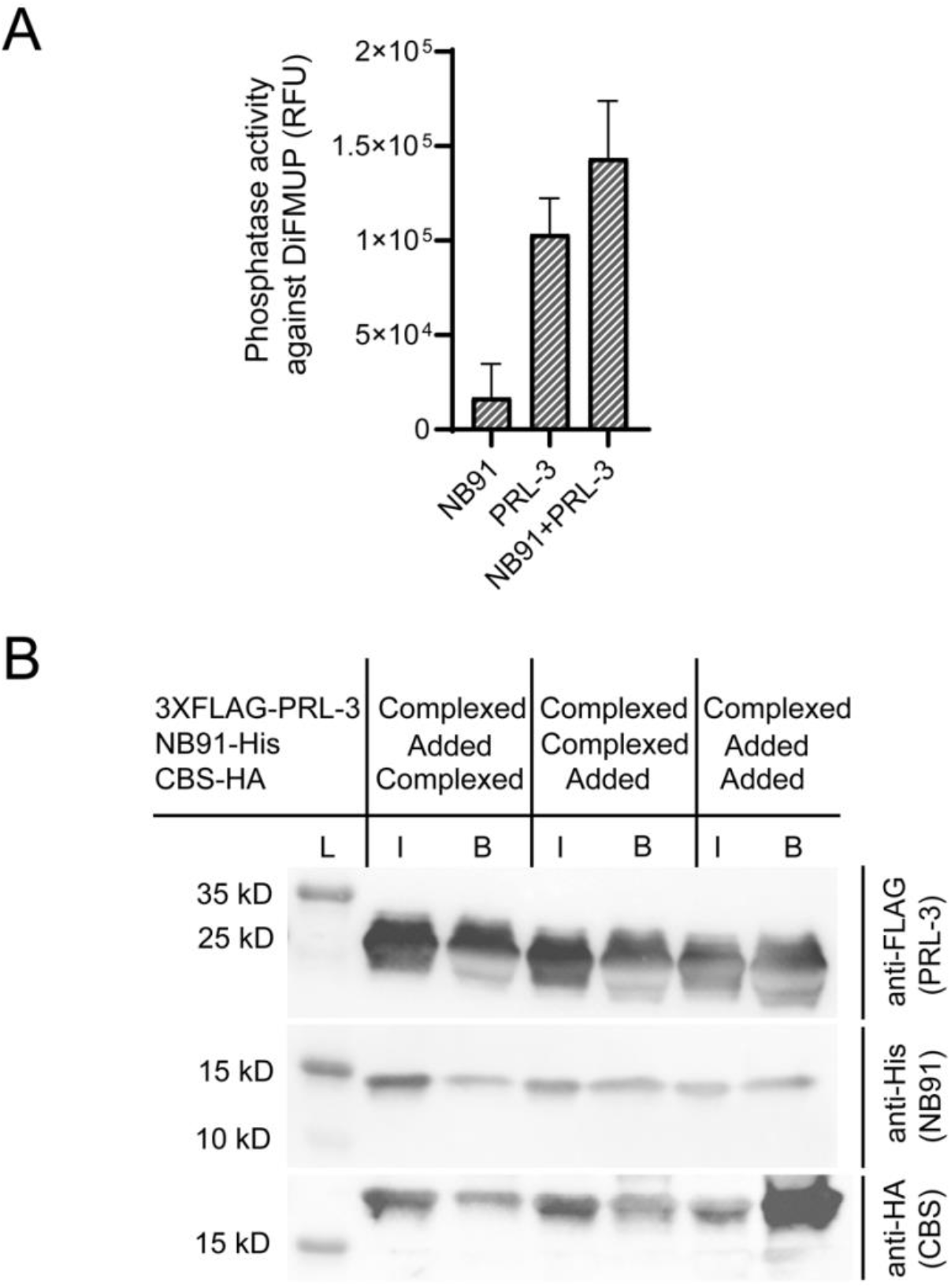
Nanobody 91 binds outside of the PRL-3 active site. (A) Phosphatase activity of each protein, alone or in complex, against a generic diFMUP substrate. DiFMUP fluoresces when phosphate is released. The graph shows relative fluorescence units, n=12 ± standard deviation. (B) Western blot of analysis of binding partners that were immunoprecipitated with PRL-3. 3XFLAG-tagged PRL-3 was complexed with anti-FLAG beads and either the HA-tagged CBS domain of CNNM3 (left column), histidine-tagged Nanobody 91 (middle column), or neither. After 1 hr incubation, NB91-His (left column), CBS-HA (middle column), or both proteins (right column) were added to the complex before immunoprecipitation. L, ladder; I, input; B, proteins bound to FLAG beads after wash. Antibodies used for western blot are shown.

The molecular weight of diFMUP is only 292; the 15 kD nanobodies could theoretically sterically hinder a protein substrate from binding into the PRL-3 active site. The magnesium transporter CNNM3 is a well-established PRL-3 binding partner and is involved in PRL-3 pseudo-phosphatase activity (43). As shown in PDB:5TSR, the cystathionine-β-synthase (CBS) domain of CNNM3 binds to the PRL-3 active site (43). We purified the CNNM3 CBS domain fused to a hemagglutinin (HA) tag and a 3XFLAG-tagged PRL-3 (Figure S4A-B) to examine the impact of nanobody binding on PRL-3:substrate interactions. Initially, we determined that recombinant 3XFLAG-PRL-3 can be pulled down by anti-FLAG beads, while Nanobody 91 and CBS-HA are only pulled down by anti-FLAG beads when complexed with 3XFLAG-PRL-3 (Figure S4C-D), indicating no background reactivity of the latter two proteins with anti-FLAG beads. We found that nanobody 91 could still be co-immunoprecipitated with PRL-3 even if PRL-3 was already in complex with CBS-HA, and that the nanobody:PRL-3 complex did not prevent CBS binding and co-immunoprecipitation (Figure 4B). Finally, we complexed 3XFLAG-PRL-3 to anti-FLAG beads, followed by incubation with both recombinant CBS-HA and Nanobody 91 simultaneously and found that neither protein outcompeted the other for binding of PRL-3, as both proteins were co-immunoprecipitated (Figure 4B). Taken together, these data indicate that anti-PRL-3 nanobodies bind away from the active site of PRL-3 and are unlikely to interfere with PRL-3 phosphatase or pseudo-phosphatase activity.

### Anti-PRL-3 nanobodies immunoprecipitated PRL-3 but not PRL-1 or PRL-2 from HEK293T overexpressing cell lysates

PRL-3 substrates and binding partners remain primarily undefined, in part due to insufficient tools for cell-based studies—the current commercially available antibodies that can recognize PRL-3 in its native state have not been extensively validated for specificity towards PRL-3 over other PRLs. We tested the ability of the anti-PRL-3 nanobodies to immunoprecipitate PRL-3 protein from cells expressing FLAG-tagged PRL-1, PRL-2, or PRL-3. All nanobodies selectively pulled-down PRL-3 over PRL-1 and PRL-2 (Figure 5A). Some nanobodies were more specific to PRL-3 than others; for example, nanobodies 4, 16, 19, and 84 pulled down small amounts of 3XFLAG-PRL-1 and PRL-2. Successful nanobody coupling to Dynabeads was confirmed by the presence of 6XHis-tag in all samples (Figure 5B). Beads alone do not immunoprecipitate 3XFLAG-PRL-3 (Figure S5), indicating that all FLAG-PRL-3 pulldown was due to the presence of the nanobody. Total protein loading is also shown in Figure S6A-H to represent loading controls for each blot. In total, anti-PRL-3 nanobodies 10, 26, and 91 can be used to specifically immunoprecipitate PRL-3, with no binding to PRL-1 or PRL-2.

**Figure 5.**
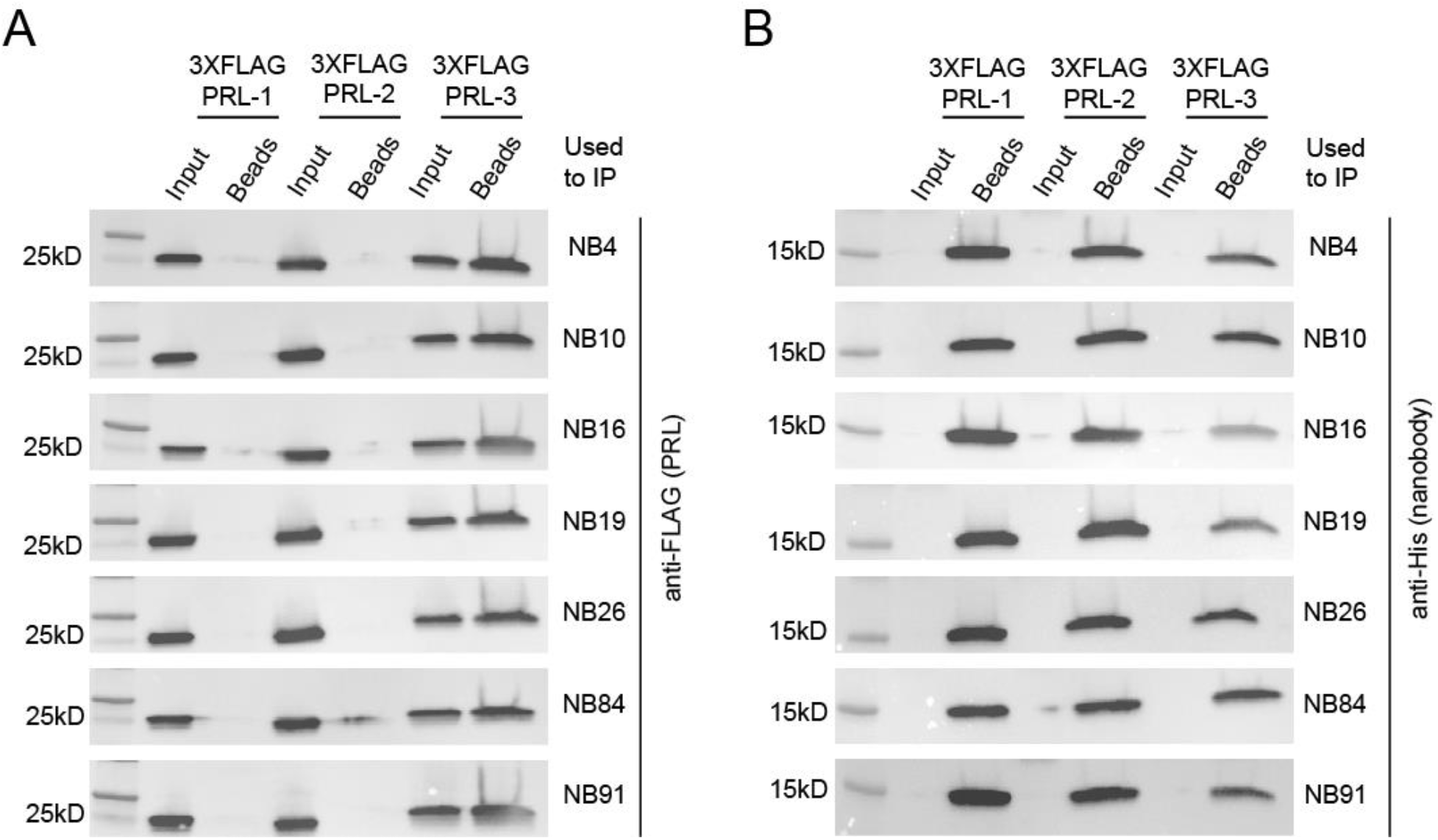
Nanobodies selectively immunoprecipitate PRL-3 from HEK293T cell lysate. PRL-3 specific nanobodies coupled to superparamagnetic Dynabeads® M-270 Epoxy beads were used in immunoprecipitation assays with lysates from HEK293T cells transduced with 3XFLAG-PRL-1, −2 or −3. (A) All nanobodies pulldown 3XFLAG-PRL-3 with minimal to no pulldown of 3XFLAG-PRL-1 or 3XFLAG-PRL-2. (B) Successful nanobody coupling to Dynabeads in all groups was verified using an antibody against the C-terminal 6XHis-tag present on each nanobody.

### Anti-PRL-3 nanobodies specifically detect PRL-3 in fixed cells in immunofluorescence assays

Our next goal was to determine if nanobodies could specifically detect PRL-3 in fixed cells via immunofluorescence, which can be used to examine PRL-3 localization and trafficking. The human colon cancer cell line HCT116 was transfected with CMV:GFP-PRL-1, −2, or −3 constructs to visualize the PRLs and determine the extent to which the anti-PRL-3 nanobodies co-localize with each of them. Nanobody 91 co-localized with GFP-PRL-3, which was found mainly at the plasma membrane and rarely in the nucleus (Figure 6), which are previously described sites of PRL-3 localization (21,23). The anti-PRL-3 nanobody did not stain cells expressing GFP-PRL-1, GFP-PRL-2, or the GFP vector control. All seven anti-PRL-3 nanobodies tested were similarly specific for PRL-3 over PRL-1 and PRL-2 (Figure S7-12), although some nanobodies exhibited a higher non-specific background signal, such as nanobody 84 (Figure S12).

**Figure 6.**
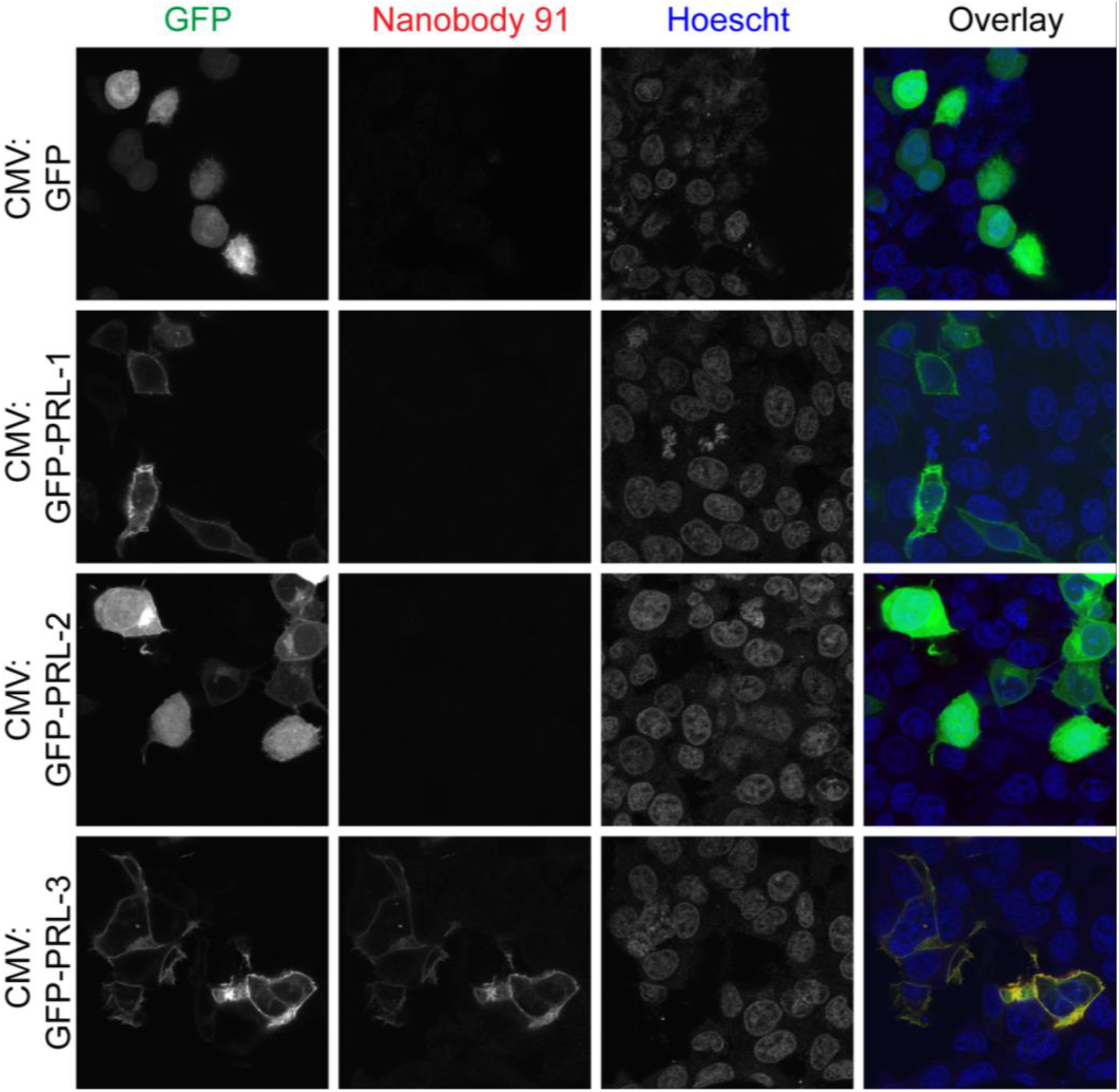
Nanobodies are specific to PRL-3 in immunofluorescence assays. HCT116 colorectal cancer cells were transfected with CMV:GFP, CMV:GFP-PRL-1, CMV:GFP-PRL-2, or CMV:GFP-PRL-3 for 24 hours prior to cell fixation and permeabilization. Immunofluorescence assays were completed with 1:100 1 mg/mL NB91 followed by 1:400 Alexa Fluor® 594-AffiniPure Goat Anti-Alpaca IgG, VHH domain, showing that nanobodies detect and co-localize with PRL-3 but not PRL-1 or PRL-2.

### An N-terminal tag impacts PRL-3 localization in HCT116 colorectal cancer cells

While exploring GFP-PRL-3 localization in the cell, we observed that this form of PRL-3 localized mainly to the cell membrane, with occasional foci present in the nucleus (Figure 6). GFP is ~28 kD in size, doubling the size of the PRL-3 that is being expressed in HCT116 cells. Researchers often assume that N- and C-terminal tags have little influence on the secondary and tertiary structures and localization of fused proteins (44). However, several reports have demonstrated that the use of GFP may impact the biological activity of fusion proteins (45–47), including cellular localization (48). Many past studies examining PRL-3 localization, which have primarily characterized PRL-3 as membrane exclusive due to the presence of a C-terminal prenylation, have utilized N-terminal tags such as Myc (20,49) or EGFP (49,50). We wanted to use our PRL-3 nanobodies to determine if tagged versions of PRL-3 localize differently than wild-type, untagged PRL-3.

We examined the localization of GFP-PRL-3 and 3XFLAG-PRL-3. FLAG tags are often utilized in immunoprecipitation experiments, as FLAG is only 3kD and made of primarily charged amino acids. We expressed GFP-tagged, 3XFLAG-tagged, and wild-type PRL-3 at equal levels in HCT116 cells (Figure 7A). Probing with Nanobody 26 revealed that tagged PRL-3 was strongly localized to the membrane, compared to the cytoplasm, with punctate staining at the nucleus (Figure 7B-C). In comparison, untagged PRL-3 is evenly distributed across the cytoplasm and cell membrane (Figure 7C). These findings also hold true when examining localization patterns with nanobodies 19 and 91 (Figure S13 and S14). In total, PRL-3 expression is much more variable than previous studies using tagged protein might indicate; these results suggest that N-terminal tags may be affecting PRL-3 localization.

**Figure 7.**
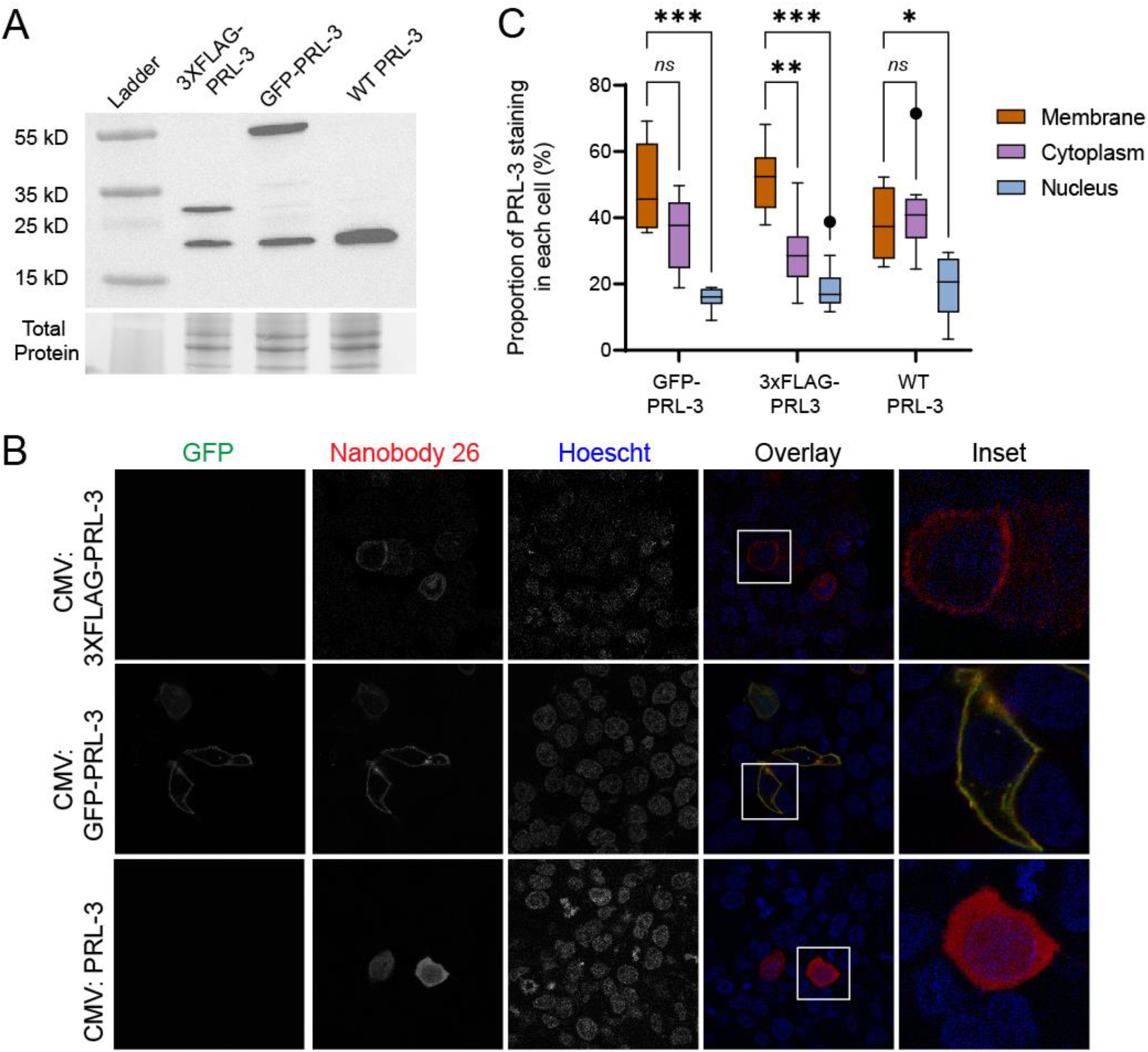
PRL-3 localization assessed by Nanobody 26 is altered by N-terminal 3XFLAG and GFP tags. (A) PRL-3 western blot indicates a similar expression of exogenous proteins. 3XFLAG-PRL-3 can be seen at ~27 kD, and GFP-PRL-3 is shown at ~55 kD. CMV-PRL-3 and endogenous PRL-3 are represented at 22 kD. (B) Immunofluorescence of HCT116 cells transfected with CMV:3XFLAG-PRL-3, CMV:GFP-PRL-3, or CMV-PRL-3, as indicated. Cells were stained with anti-PRL-3 nanobody 26 followed by an anti-alpaca VHH coupled to Alexa594 secondary antibody for visualization. (C) ImageJ quantification of nanobody/PRL-3 staining. Groups were compared using a Two-Way Anova with Tukey’s Multiple Comparisons Test where \**p* < 0.05, \*\**p* < 0.01, \*\*\**p* < 0.001.

## Discussion

The PRL family of proteins has emerged as important in cancer progression, with PRL-3 now recognized as a bona fide oncogene. However, the mechanisms by which PRL-3 promotes tumor growth and spread are largely unknown and are essential to define before PRL-3 inhibitors can be used in the clinic. A significant roadblock in understanding the role of PRL-3 in cancer is a lack of tools to study this protein. While PRL antibodies and inhibitors have been used for research purposes, they often come with caveats. For example, the allosteric PRL-3 inhibitor JMS-053 equally targets the entire PRL family, PRL-1, −2, and −3 (27). One of the commercially available antibodies often used in the literature (R&D Systems MAB3219) has been validated as specific towards PRL-3 yet was generated against denatured protein and is only useful for immunoblot. PRL-3-zumab requires an *in vivo* microenvironment for its anti-cancer activity and has not been made widely available (30,51).

Much like PRL-3-zumab, the anti-PRL-3 nanobodies we have generated are specific to PRL-3 over PRL-1 and PRL-2, with a PRL-3 binding affinity in the low nM range. Nanobodies carry advantages over their conventional antibody counterparts in general and in terms of PRL-3. So far, PRL-3-zumab has been applied as an extracellular reagent and is hypothesized to bind PRL-3 on the cell surface to induce an immune response to kill cancer cells (30). While nanobodies are 10-fold smaller compared to conventional antibodies and are stable proteins, they are unlikely to penetrate cell membranes via passive diffusion. Researchers have recently focused on developing cell-penetrating nanobodies. For example, Herce et al. attached intracellularly stable cyclic arginine-rich cell-penetrating peptides (CPPs) to nanobodies (52). CPPs increase the size of cargo that can be delivered efficiently into living cells (53). This strategy was successful in delivering nanobodies targeted against GFP (27.5 kD), GFP–PCNA (63 kD), and the therapeutically relevant Mecp2– GFP fusion protein (83 kD) into HeLa cells to study protein interactions (52). Similar technologies could be used to deliver anti-PRL-3 nanobodies for studies on PRL-3 localization, trafficking, or function in living cells. Nanobodies have also emerged as a valuable tool to neutralize target proteins involved in disease, including SARS-CoV-2 (54)—delivery of anti-PRL-3 nanobodies into the cell would position it as a useful biological therapeutic in a variety of cancers.

A major goal in the PRL field is to complete in-depth, high-resolution structural studies of PRL-3, especially in complex with current inhibitors or substrates. This information can give insight into designing better small molecules to target PRL-3 and identify novel sites to target the protein. However, PRL-3 has been difficult to crystalize without a substrate-bound in its active site, as it can move between open and closed conformation (24). Current PRL-3 antibodies are not practical to stabilize PRL-3 for crystallization, as they are large, glycosylated, multi-domain proteins, which are not suitable for applications such as X-ray crystallography. The small size, high stability, and high specificity of nanobodies lend them well towards acting as chaperones in structural studies (55). Our data demonstrate that anti-PRL-3 nanobodies interact with PRL-3 in solution and bind with a high affinity outside PRL-3’s active site, based on HDX-MS parameters. Therefore, the nanobodies may help stabilize the PRLs in a single conformation for crystallization studies. A high-resolution crystal structure of PRL-3 with an unbound active site would be useful in *in silico* drug design and substrate identification.

Finally, because our nanobodies bind outside of PRL-3’s active site, they may be useful in identifying PRL-3 substrates that are important in cancer progression. We found that nanobodies can be used in co-immunoprecipitation and immunofluorescence without occluding any PRL-3 binding partners. Importantly, we observed PRL-3 localization could significantly change with the addition of an N-terminal tag. Both FLAG and GFP-tagged PRL-3 localized more often to the membrane than wild-type PRL-3, which was found throughout the cell. One of the current aspects of our nanobodies that must be addressed is that all experiments in this study utilized overexpressed PRL-3. Further optimization of these assays is necessary so that PRL-3 nanobodies can be utilized to study and target endogenous PRL-3. We demonstrated the wide variety of applications for the PRL-3 nanobodies and showed they bind cellularly expressed PRL-3; we expect that nanobodies will bind endogenous PRL-3 as well.

PRL-3’s function may vary based on cellular location. For example, PRL-3 binds CNNM3 at the membrane to regulate magnesium transport via pseudo-phosphatase activity, but PRL-3 is also established to dephosphorylate several different cytoplasmic proteins. How different functions of PRL-3 might contribute to different aspects of cancer progression is currently unknown. Much research related to PRL-3 has used N-terminus tagged protein due to a lack of antibody tools. We hope that the availability of nanobodies that can specifically bind to untagged PRL-3 will lead to new discoveries regarding PRL-3 function.

In summary, we have developed the first alpaca-derived single domain antibodies against PRL-3 and showed that they could specificity detect PRL-3 in multiple *in vitro* assays, in human cell lysates overexpressing PRL-3, and *in situ* in fixed cancer cells. At the same time, they do not interfere with PRL-3 phosphatase activity or substrate binding. These nanobodies have begun to fill an important gap in the tools needed to study PRL-3 function in normal physiology and cancer and have the potential to provide valuable insight into PRL-3 substrates, trafficking, structure, and inhibition.

## Experimental Procedures

### Plasmids and other reagents

To generate recombinant protein for alpaca immunization, human PRL-3 cDNA was amplified with gene-specific primers and cloned into the bacterial expression vector, pSKB3. pSKB3 is a modified pET-28b vector, where a thrombin cleavage site was replaced by a TEV protease cleavage site and was originally constructed by Dr. Steve Burley, Rutgers, The State University of New Jersey. pSKB3 was used to clone the recombinant PRL-1, −2, and −3 at NheI and XhoI restriction sites using T4 ligase.

The 3XFLAG-tagged PRL mammalian expression plasmids were made by cloning full-length PRL-1, −2, or −3 human cDNA into a p3XFLAG-CMV-14 expression vector (Sigma, E7908). Then 3XFLAG-PRLs were cloned into pLenti-CMV-puro (Addgene 17452) to make plenti-CMV-3XFLAG-PRL-puro constructs.

The 3XFLAG-tagged PRL-3 protein expression vector was generated by amplifying 3XFLAG-PRL-3 from a pCMV-3XFLAG-PRL-3 vector via PCR and cloned into pSKB3 utilizing NheI and XhoI restriction sites using T4 ligase. The CBS-HA protein expression vector was made by cloning full-length CNNM3 CBS domain pair gBlocks™ Gene Fragments (IDT) into pSKB3 using NheI and XhoI restriction enzyme sites and T4 ligase.

The GFP-tagged and untagged PRL overexpressing plasmids were made by cloning full-length PRL-1, −2, or −3 gBlocks™ Gene Fragments (IDT) into the pcDNA™3.1 (-) (Invitrogen V79520) at BamHI and HindIII restriction sites. A GFP gBlock was subsequently cloned into each of the pCDNA3.1-PRL plasmids to generate CMV:GFP-PRL fusion constructs at NotI and BamHI restriction sites.

### Production, panning, and sequencing of nanobodies

Nanobodies were produced by the University of Kentucky Protein Core, as previously described (55). Briefly, 100 μg of recombinant PRL-3 antigen (See Protein purification) was subcutaneously injected into alpacas once per week for six weeks to boost nanobody presence in the immune system. 3-5 days following the final injection 50 mL of alpaca blood was harvested to isolate peripheral blood lymphocytes by density gradient centrifugation. RNA was isolated from these lymphocytes, and cDNA was synthesized using reverse transcriptase. A bacteriophage display cDNA library was made by cloning potential VHH regions, with restriction enzymes, into the phage display vector pMES4. pMES4 phage was expressed with the VHH insert fused to gene III of the filamentous phage for the production of the phage solution. Two rounds of phage display against PRL-3, as demonstrated previously (55), utilizing this cDNA library yielded 32 potentially VHH positive clones. Positive clones were confirmed using a VHH specific primer pEX-Rev (CAGGCTTTACACTTTATGCTTCCGGC) and sequencing by Eurofins Genomics. DNA sequences were translated using the ExPASy Bioinformatics Resource Portal Translate Tool (https://web.expasy.org/translate/), where they were analyzed for nanobody components including pelB sequence and 6XHis-tag followed by a stop codon. 16 of 32 clones embodied these components and were carried through to following experiments.

### Cell lines and cell culture

Both of the human cell lines used in this study (HEK293T, HCT116) were authenticated by short tandem repeat (STR) profiling and tested for mycoplasma contamination prior to experiments using the LookOut® Mycoplasma PCR Detection Kit (Sigma, MP0035-1KT). HEK293T (ATCC CRL-3216) and HCT116 (ATCC CCL-247) cells were grown in 1X DMEM (Thermofisher, 11965092). For all, media were supplemented with 10% heat-inactivated fetal bovine serum (R&D Systems, S11150H, Lot. H19109). Cells were cultured at 37 °C with 5% CO_2_. To overexpress the CMV:PRL-3, CMV:GFP-PRL, and CMV:3XFLAG-PRL plasmids, cells were transfected using Lipofectamine 3000 (Thermofisher, L3000-015) following the manufacturer’s protocol. HEK293T stably expressing PRL cell lines were selected with 1 μg/mL puromycin (Thermofisher, A1113803).

### Protein purification

pSKB3-PRL, pSKB3-CBS-HA, pSKB3-3XFLAG-PRL-3, and pMES4-Nanobody expression plasmids described previously were transformed into and expressed using the One-Shot BL21 Star DE3 bacterial cell line (Invitrogen, C601003) by stimulating induction with 0.5 mM IPTG (Fisher Scientific, BP175510) for 16 hours at 16°C following a culture O.D._600_ of 0.6. Cells were pelleted at 5,000 rpm for 15 minutes at 4°C and resuspended in 10 mL of lysis buffer [300 mM NaCl (VWR BDH9286), 20mM Tris pH 7.5, 10 mM Imidazole pH 8.0 (Sigma-Aldrich I2399), 1:1000 protease inhibitor cocktail (Sigma-Aldrich P8465)] per gram of cell pellet and lysed using a microfluidizer (Avestin, EmulsiFlex-C5). Debris was pelleted at 18,000 rpm for 50 minutes at 4°C, and the lysate was run over 1 mL columns (Biorad, 7321010) packed with Ni-NTA Resin (VWR, 786-940). PRLs (Figure S15) and CBS (Figure S6A) were eluted with 2 mL of elution buffer (300 mM NaCl, 20 mM Tris pH 7.5, and 250 mM Imidazole pH 8.0). Nanobodies underwent two elution steps, the first with 30 mM Imidazole elution buffer and the second with 250 mM Imidazole elution buffer (Figure S16). The N-terminal 6XHis-tag on recombinant PRLs was cleaved using TEV protease (gift from Dr. Konstantin Korotkov), and samples were reapplied to the Ni-NTA column to remove uncleaved protein. Recombinant nanobodies remained with their C-terminal 6XHis-tag intact. All samples underwent buffer exchange to remove imidazole (300 mM NaCl, 20 mM Tris pH 7.5), and PRLs and nanobodies were further purified using a Superdex 200 Increase 10/300 GL column (GE, 28990944) on an ÄKTA purification system in buffer containing 100 mM NaCl and 20 mM HEPES (Fisher Scientific, BP310-100) pH 7.5. Purification was verified by running samples on 4-20% Mini-PROTEAN TGX Stain-Free Gels (Biorad 4568094) (Figures S1D and S2B). The purest fractions were pooled, concentrated together, flash-frozen on dry ice, and stored at −80°C.

### ELISA for Nanobody/PRL binding specificity

Recombinant, purified, PRL-1, −2, and −3 were plated at 1 μg/mL (100 μl) in sodium bicarbonate buffer [0.42g sodium bicarbonate (Fisher Scientific, BP328-500) in 50 mL diH_2_0] in Corning® 96 Well EIA/RIA Assay Microplates (Sigma, CLS3590) and incubated for 16-20 hours at 4°C. Plates were washed three times with 0.05% PBST and loaded with a blocking solution of 0.5% BSA (Fisher Scientific, BP9706100) in 0.1% PBST for 1 hour at room temperature. Blocking buffer was removed, and nanobodies were diluted to 1 μg/mL, or designated concentration for dosing experiments, and incubated in wells for 1 hour at room temperature. Wells were washed 3 times in PBS and incubated with 1:1000 anti-His HRP antibody (GenScript, A00612, Lot. 19K001984) for 1 hour at room temperature. Plates were washed 3 times with PBS and developed with TMB 2-Component Microwell Peroxidase Substrate Kit (Seracare, 5120-0053). Reactions were stopped after 90 seconds with 0.1 N HCl (Fisher Scientific, A144500) and read on a Biotek Synergy Multi-mode Plate Reader at 450 nm. Controls included PRL only wells to specify lack of a 6X-His-tag, nanobody, and secondary only wells to specify the necessity of PRL presence for binding, and buffer only to provide evidence that sodium bicarbonate and BSA did not elucidate a colorimetric change. Raw data from all control wells were pooled for each plate, and experimental wells were normalized to controls by dividing individual wells by average control wells. Individual well readouts were then placed in Prism 7 in a Grouped format Table, where values for two replicate experiments were graphed for relative absorbance at 450 nm compared to the average of control wells.

### Immunoprecipitation of PRL-3 with Nanobody Coupled Dynabeads

PRL-3 nanobodies were coupled to Dynabeads (Life Technologies, 14311D) for downstream 3XFLAG-PRL immunoprecipitation following the manufacturer’s instructions. HEK293T cells (~20 million) stably expressing pLenti-CMV-puro (Addgene 17452) empty vector, PRL-1, PRL-2, or PRL-3 under 1 μg/mL puromycin selection were lysed for 30 minutes with intermittent vortexing in Pierce IP lysis buffer (Thermo 87788) supplemented with 1% protease inhibitor cocktail (IP buffer) at 500 μl per 10 million cells and spun at 12,000 rpm for 10 minutes at 4°C to pellet cell debris. Protein concentration was quantified using the QuickStart Bradford 1X Dye Reagent (Biorad, 5000205). 150 μL of nanobody-coupled beads were washed in 1 mL of PBS for 5 minutes, then equilibrated in 500 μL of IP buffer for 5 minutes. 2.5 mg of total extracted protein was added to the equilibrated nanobody-beads complex for incubation at 4°C overnight with rocking. After washing the beads-protein complex in cold PBS four times, 50 μL 2x Laemmli Sample Buffer (Biorad, 161-0737) with 2-Mercaptoethanol (Fisher Scientific, 03446I-100) was added to the beads, the mixture was boiled at 95°C for 10 minutes, and the supernatant was collected for western blot analysis.

### Co-Immunoprecipitation of CBS-HA and Nanobody 91 with 3XFLAG-PRL-3

Anti-FLAG M2 magnetic beads (Sigma, M8823-5ML, Lot. SLCF4223) were prepared by washing in No Imidazole Buffer (300 mM NaCl, 20 mM Tris pH 7.5) twice. Initial complexing to beads occurred for 1 hour at 4°C, with secondary complexing occurring for a second hour at 4°C. CBS-HA and Nanobody 91 were both added in 1:1.1 molar ratios to 3XFLAG-PRL-3 complexed beads. Following the second complexing step, the supernatant was removed, and beads were washed four times with No Imidazole Buffer. Beads were eluted with 50 μL 2x Laemmli Sample Buffer (Biorad, 161-0737) with 2-Mercaptoethanol (Fisher Scientific, 03446I-100). The eluates were boiled at 95°C for 10 minutes, and the supernatant was collected for western blot analysis.

### Western Blot

30 μg of total protein for input or 45 μl of pulldown supernatant was loaded into 4–20% Mini-PROTEAN® TGX Stain-Free™ Protein Gels. Total protein was assessed through stain-free imaging on a Biorad ChemiTouch Imaging System, which allows the use of total protein as the loading control. Protein was transferred onto the PVDF membrane (Biorad, 162-0255) using the Trans-Blot Turbo Transfer System (Biorad 1704150). Membranes were blocked with 5% milk in 0.1% TBST for 1 hour and probed with one of the following antibodies at the designated dilution overnight at 4°C. 1:3000 Monoclonal ANTI-FLAG® M2 antibody (Sigma, F1804, Lot. SLBK1346V), 1:1000 anti-His HRP antibody (GenScript, A00612, Lot. 19K001984), 1:1000 anti-HA-Tag (C29F4) Rabbit mAb (Cell Signaling, 37245, Lot. 8), or 1:1000 anti-Human/Mouse/Rat PRL-3 Antibody (R&D Systems, MAB3219, Lot. WXH0419091). Following three washes with 0.1% TBST, secondary HRP-conjugated 1:2500 anti-mouse IgG antibody (Cell Signaling, 7076S, Lot. 33) or 1:5000 anti-Rabbit IgG HRP Linked F(ab′)2 (Sigma, NA9340V, Lot. 17065618) was added for 1 hour and membranes were imaged using Clarity Western ECL Substrate (Biorad, 1705061).

### Immunofluorescence in fixed PRL overexpressing cells with nanobodies and quantification with ImageJ

HCT116 cells were plated at 5,000 cells per well in 96-well black glass-bottomed plates (Cellvis, P96-1.5H-N) and transfected with either pcDNA3.1-PRLs, GFP-PRLs, or pLV-3XFLAG-PRLs (Addgene 123223) as previously described in the Cell lines and cell culture section. All solution exchanges and imaging occurred in the 96-well plate. 24 hours post-transfection cells were fixed in 4% paraformaldehyde (VWR, AAJ61899-AK) for 15 minutes, rinsed in PBS, permeabilized for 10 minutes in 1% Triton X-100 (Sigma, X100-100), and rinsed in PBS. A blocking solution of 2% BSA in PBS was applied for 1 hour to all wells. All nanobodies were diluted to 1 mg/ml in blocking solution, further diluted 1:100, and incubated with the fixed cells for 1 hour at room temperature followed by five PBS washes. Detection was carried out using an anti-alpaca IgG VHH conjugated to Alexa Fluor-594 (Jackson ImmunoResearch, 128-585-232) diluted 1:400 in blocking solution and counterstained with Hoechst, 1:1000 dilution (ThermoFisher, H3570). All wells were washed in PBS five times prior to imaging. Images were acquired at the University of Kentucky Light Microscopy Core using a Nikon A1R confocal microscope using a 40X water objective. Images were processed in Adobe Photoshop 2020 to both increase image brightness and overlay the 405 (Hoescht), 488 (GFP-PRL-3), and 561 (Nanobodies) channels. Channels were pseudocolored by RGB channels. To quantify the average grey area in three cellular compartments (membrane, cytoplasm, and nucleus), we utilized Image J. Briefly, Nikon files (.nd2) were converted to .tif files, and images from Channel 3 (561nm) were opened in Adobe Photoshop 2022 where brightness and contrast were adjusted to match between the three different tags. Adobe Photoshop 2022 images were saved as .tif files and opened in ImageJ. The Straight tool, or the line tool, was selected from the toolbar to draw and measure a line across a single cell. Five 120 pixel rectangles were drawn to on a single cell, placed on the two points on the line the showed the plasma membrane, two sides of the cytoplasm, and one to denote the nucleus. The average grey value was measured for each rectangle was measured, and the plasma membrane and cytoplasm results for averaged for each cell. For each condition (3XFLAG, GFP, or WT), ten cells were measured. From the toolbar, the Analyze menu was selected, and then Plot Profile was selected. Average grey value quantifications were exported to Microsoft Excel as .xls files and graphed using Graphpad Prism 9. Statistical analysis was done using a two-way ANOVA with multiple comparisons to examine the localization of three different types of PRL-3 to three different cellular compartments.

### Phosphatase Assay

2.5 μM of recombinant PRL-1, −2, or −3 was mixed with 2.5 μM of each nanobody in black 384-well plates (Thermo Scientific, 164564) and incubated at room temperature for 1 hour in Reaction Buffer (20 mM Tris, 150 mM NaCl). Following incubation, the recombinant protein mixtures were combined with 12.5 μM diFMUP (Life Technologies, E12020), added to 384-well plates, and incubated for 20 minutes in the dark at room temperature. Fluorescence intensities were measured on a Biotek Synergy Multi-mode Plate Reader at 360 nm/460 nm excitation and emission wavelengths, respectively. Raw values for non-substrate-containing controls were averaged and subtracted from values of wells incubated with the substrate to remove background fluorescence. Raw values were transferred to Prism 7 software in a Grouped format, where two replicate experiments were combined for final data processing.

### BLItz technology for K_D_ determination

Anti-Penta-HIS (HIS1K) Biosensors (ForteBio, 18-5120) were hydrated for 10 minutes in 1X kinetics buffer (ForteBio, 18-1105, Lot. 20070082) in a 96-well black bottom plate. The Advanced Kinetics Assay protocol on BLItzPro1.3 software was used, with five steps in the following order: Initial Baseline 30 seconds, 250 μl 1X kinetics buffer (ForteBio, 18-1105); Loading 30 seconds − 4 μl of ligand (10 μg/mL nanobody); Baseline 30 seconds, 250μl 1X kinetics buffer; Association 5 minutes, 4 μl analyte (PRL-3); Dissociation 2 minutes, 250 μl 1X kinetics buffer. Global K_D_ for each nanobody was analyzed with BLItzPro 1.3 software using baseline correction for both the association and dissociation steps.

### Hydrogen Deuterium Exchange Mass Spectrometry

The coverage map for PRL-3 was obtained from undeuterated controls as follows: 3μL of 20μM apo PRL-3 in 20 mM HEPES pH 7.5, 100 mM NaCl was added to 9 μL of buffer (20 mM HEPES, pH 7.5, 100 mM NaCl). The reaction was quenched with 14 μL of ice-cold quench (100 mM glycine, pH 2.5, 7 M Guanidine HCl, 10 mM TCEP) then diluted with 54 μL of ice-cold dilution buffer (100 mM Glycine, pH 2.5, 10 mM TCEP). 40 μL of this solution was injected into a Waters HDX nanoAcquity UPLC (Waters, Milford, MA) with in-line protease XIII/pepsin digestion (NovaBioAssays). Peptic fragments were trapped on an Acquity UPLC BEH C18 peptide trap and separated on an Acquity UPLC BEH C18 column. A 7 min, 5–35% acetonitrile (0.1% formic acid) gradient was used to elute peptides directly into a Waters Synapt G2-Si mass spectrometer (Waters, Milford, MA). MSE data were acquired with a 20–30 V ramp trap CE for high energy acquisition of product ions as well as continuous lock mass (Leu-Enk) for mass accuracy correction. Peptides were identified using the ProteinLynx Global Server 3.0.3 (PLGS) from Waters. Further filtering of 0.3 fragments per residue was applied in DynamX 3.0.

For each complex and the apo PRL-3, the HD exchange reactions and controls were acquired using a LEAP autosampler controlled by Chronos software. The reactions were performed as follows: 1.5μL of 40μM PRL-3 in buffer with 1.5 μL 100 μM of the given nanobody in buffer (or 1.5 μL of buffer for the apo PRL-3 experiments) was labeled with 9 μL of 20 mM HEPES, pD 7.5 ^2^H_2_O based buffer with 100 mM NaCl, and quenched with 14 μL of ice-cold quench (100 mM Glycine, pH 2.5, 7 M Guanidine HCl, 10 mM TCEP) then diluted with 54 μL ice-cold dilution buffer (100 mM Glycine, pH 2.5, 10 mM TCEP). 40 μL of this solution was used for analysis. All deuteration time points were acquired in triplicates. Back exchange correction was performed against fully deuterated controls. Fully deuterated controls were done by incubating 2μL of 40 μM PRL-3 in 8 μL of unfolding buffer (100mM Glycine pH 2.5, 7 M Guanidine HCl, 10 mM TCEP) for two hours. Then 3 μL of the unfolded solution was deuterated for 2 hours in 9 μL of a deuterium-based buffer. This was quenched with 14 μL of ice-cold quench (100 mM Glycine, pH 2.5, 10 mM TCEP) then diluted with 54 μL ice-cold dilution buffer (100 mM Glycine, pH 2.5, 10 mM TCEP). 40 μL of this solution was used for analysis. The deuterium uptake for all identified peptides with increasing deuteration time and for the fully deuterated control was determined using Water’s DynamX 3.0 software.

The normalized percentage of deuterium uptake (%D_t_) at an incubation time t for a given peptide is calculated as such:

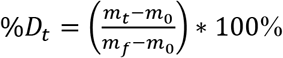

with m_t_ the centroid mass at incubation time t, m_0_ the centroid mass of the undeuterated control, and m_f_ the centroid mass of the fully deuterated control. Percent deuteration difference was calculated as such: Δ%Dt(PRL-3 – PRL-3 NB Complex). Δ%D_t_ for each complex at each time point was then mapped onto the PRL-3 structure. Kinetic uptake plots and corresponding heatmaps were generated using an in-house python script.

## Data Availability

All data are contained in this manuscript.

## Supporting Information

This article has the following supporting information.

Table S1. Raw data for Nanobody KD calculations using biolayer interferometry.,

Figure S1. Nanobody 19 stabilizes PRL-3 structure at two sites and destabilizes PRL-3 at one interaction point.,

Figure S2. Nanobody 26 stabilizes PRL-3 structure at two sites and destabilizes PRL-3 at one interaction point.,

Figure S3. Nanobodies do not specifically inhibit the phosphatase activity of PRL-3.,

Figure S4. Protein purification and immunoprecipitation controls for detecting 3XFLAG-PRL-3, CBS-HA, and NB91-His interactions.,

Figure S5. 3XFLAG-PRL immunoprecipitation beads only controls in HEK293T cells.,

Figure S6. Total protein for HEK293T 3XFLAG-PRL IPs.,

Figure S7. Nanobody 4 can detect overexpressed PRL-3 in fixed cells.,

Figure S8. Nanobody 10 can detect overexpressed PRL-3 in fixed cells.,

Figure S9. Nanobody 16 can detect overexpressed PRL-3 in fixed cells.,

Figure S10. Nanobody 19 can detect overexpressed PRL-3 in fixed cells.,

Figure S11. Nanobody 91 can detect overexpressed PRL-3 in fixed cells.,

Figure S12. Nanobody 84 can detect overexpressed PRL-3 in fixed cells.,

Figure S13. PRL-3 is differentially localized in colorectal cancer cells when tagged versus untagged using Nanobody 19.,

Figure S14. PRL-3 is differentially localized in colorectal cancer cells when tagged versus untagged using Nanobody 26.,

Figure S15. Representative recombinant PRL protein purification.,

Figure S16. Representative recombinant nanobody protein purification.

## Acknowledgments

We would like to thank Dr. Craig Vander Kooi and Savita Sharma for supervising phage display techniques, Dr. Yelena Chernyavskaya for providing oversight in immunofluorescence experiments, and Dr. Min Wei for help with immunoprecipitation experiments. We would also like to thank the University of Kentucky Protein Core in the Center for Molecular Medicine for alpaca injections and cDNA library construction and the University of Kentucky Light Microscopy Core for aiding in confocal training.

## Funding and Additional Information

NCI R37CA227656 and NIH DP2CA228043 to J.S.B., GMaP Region 1 North Stimulus Award to C.N.S. (Research reported in this manuscript was supported by the National Cancer Institute of the National Institutes of Health under Award Number P30CA177558. The content is solely the responsibility of the authors and does not necessarily represent the official view of the National Institutes of Health).

